# Attentional dynamics of efficient codes

**DOI:** 10.1101/2021.03.29.437459

**Authors:** Wiktor Młynarski, Gašper Tkačik

## Abstract

Top-down attention is hypothesized to dynamically allocate limited neural resources to task-relevant computations. According to this view, sensory neurons are driven not only by stimuli but also by feedback signals from higher brain areas that adapt the sensory code to the goals of the organism and its belief about the state of the environment. Here we formalize this view by optimizing a model of population coding in the visual cortex for maximally accurate perceptual inference at minimal activity cost. The resulting optimality predictions reproduce measured properties of attentional modulation in the visual system and generate novel hypotheses about the functional role of top-down feedback, response variability, and noise correlations. Our results suggest that a range of seemingly disparate attentional phenomena can be derived from a general theory combining probabilistic inference with efficient coding in a dynamic environment.

## Introduction

Activity of neurons in the visual cortex is driven jointly by external stimuli and internal feedback signals from higher brain areas [1, 2]. One hypothesis suggests that such top-down modulation could adapt sensory codes to the goals of an organism and its belief about the state of the environment [2, 3]. As a result, finite neural resources could be allocated flexibly, to sharpen representations of task-relevant stimuli at the expense of task-irrelevant input. This process, known as top-down attention [1], has been the subject of experimental and theoretical research since the early days of perceptual and neural science [4]. Yet despite the extensive research, answers to several broad questions related to top-down attention remain elusive.

The first question concerns the diversity of attentional processes. Depending on the task at hand, attention prioritizes different features of the sensory input. As a consequence, attentional processes are traditionally categorized by the relevant properties of the stimulus or the environment into, e.g., object-based attention [5–7], spatial attention [8–10], or feature-based attention [11–13]. It is unclear whether these different “kinds” of attention really reflect distinct neural processes, or whether they can be multiplexed in the same neural architecture and be governed by the same set of principles.

The second question concerns the functional effects of attentional state on neural activity. Selective attention has been shown to modulate tuning curves [14], receptive fields [15], and the firing rate of individual neurons [16]. Attentional signals can also modulate the collective structure of the population activity, in particular, correlations between pairs of neurons [17, 18]. Furthermore, fluctuations in the internal state of the brain are known to dynamically affect neural firing in the visual system [19–21]. Despite the wealth of empirical observations and qualitative hypotheses, we still lack a quantitative framework for attentional modulation that is applicable to neural activity on the individual as well as the population level.

The last question concerns the normative account of top-down attentional processing. Feedback connections have been shown to aid processing in models of the visual system by enhancing desired features of the stimulus [22, 23]. In hierarchical probabilistic models of perception, attention-like modulation can aid Bayesian inference [24]. Irrespective of the model setup, existing approaches postulate attentional dynamics rather then derive them from a normative theory of neural computation [25]. Moreover, the majority of current normative theories are themselves limited either to task-independent scenarios [26, 27] or to static environments [28–30]. It is therefore unclear whether a normative theory of task-relevant sensory coding in natural and dynamically changing environments could predict attentional phenomena *ab initio*.

Here we start by addressing the last question. Our key insight places attentional processing at the intersection of two established normative theories of neural computation: probabilistic inference and efficient coding. Probabilistic inference specifies how task-relevant environmental states can be optimally estimated from unreliable sensory signals. Efficient coding specifies how finite neural resources should be allocated to encode these signals. A recently proposed fusion of these two theories [31] provides a natural framework to study attention: a process whose presumed purpose is to allocate finite resources to extract task-relevant aspects of the environment.

Building on these principles, we develop a statistical model of adaptive sensory representations in the visual cortex. The model is optimized to infer the state of a changing environment from dynamic sequences of natural images. To navigate the efficiency constraints on the total amount of neural activity, the model utilizes top-down feedback to dynamically adapt the activation thresholds of individual neurons in the sensory population. We demonstrate that key attentional processes such as object, feature, and spatial attention emerge from the same design principle: maximization of inference accuracy at minimal neural activity cost. By optimizing this single objective, the model reproduces known static and dynamic properties of neural coding in the visual cortex, and generates novel testable predictions about neural correlations and the impact of perceptual uncertainty on the population code. Taken together, our results provide a unified normative account of the dynamic, attentionally-modulated sensory representations in the visual cortex.

## Results

We consider a scenario depicted in Fig. 1, where the aim of the sensory system is to keep track of a changing latent state of the environment. This latent state, denoted by 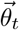 and evolving in time *t*, might correspond to a behaviorally relevant quantity, such as the position of a moving target. The brain does not have direct access to this latent state, and has to infer it from a stream of high-dimensional stimuli 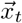. The stimuli are encoded by a resource-constrained population of sensory neurons whose instantaneous responses are denoted by 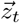. A sensory representation of the current stimulus is conveyed via feed-forward connections to a brain region which performs a specific inference (a perceptual observer). To solve this inference optimally, the observer combines the stimulus representation 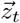 with its internal model of the world into a posterior distribution over the current state of the environment 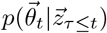. The posterior distribution is used to extract a point-estimate of the state of the environment 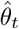, and the predicted future distribution of stimuli, which we denote as 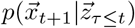. Based on this prediction, optimal parameters for the sensory population are computed and conveyed back upstream, via feedback connections. In that way, the sensory population can use its finite resources to retain only those features of the stimulus which are relevant to the perceptual observer at any given moment [31].

**Figure 1:**
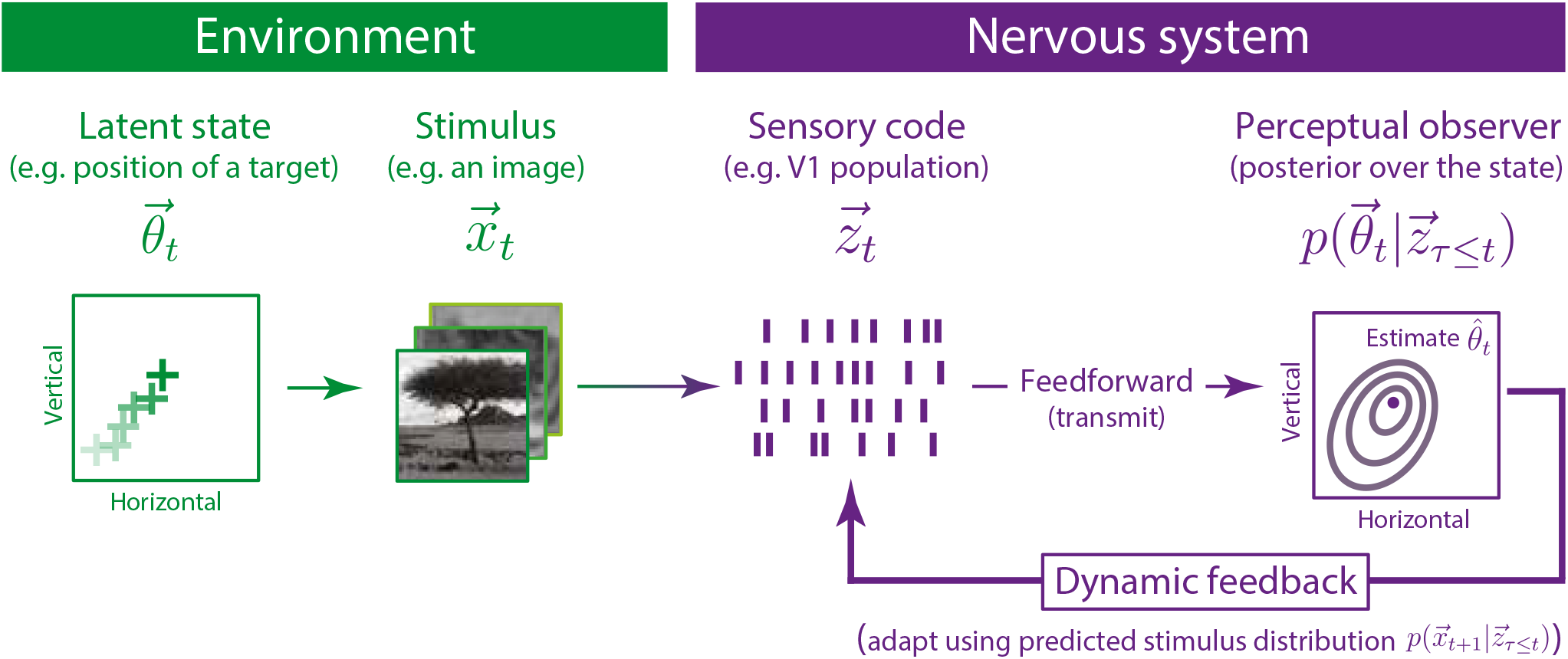
Adaptation of the sensory code for perceptual inference in a dynamic environment. Continually evolving state of the environment 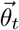 gives rise to a sequence of stimuli 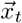, which are encoded by a population of sensory neurons into neural responses 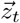. The properties of sensory neurons (e.g., their gain, receptive fields, recurrent interactions) are not fixed, but can be adapted moment-by-moment via feedback connections from higher brain areas. The normative approach we study here considers a scenario where sensory neurons optimally adapt their activation thresholds, leading to maximally accurate inference of the state of the environment by the perceptual observer, at minimal activity cost in the sensory population.

### Model of adaptive coding in the visual cortex

Following the rationale of Fig. 1, we develop a model of adaptive coding in the visual cortex (Fig. 2A, B) which is an extension of the well-known sparse coding model of V1 [26]. In the sparse coding model, a population of sensory neurons, each encoding a single image feature, forms a distributed representation of natural images. Preferred features of individual neurons are optimized to reconstruct natural images with minimal error, while maximizing the sparsity of neural responses (see Methods). The resulting features resemble receptive fields of V1 neurons and can be conveniently visualized for the entire population [27] (Fig. 2C). While sparse encoding is highly nonlinear and requires inhibitory interactions between the neurons [32], images can be linearly decoded from the population activity.

**Figure 2:**
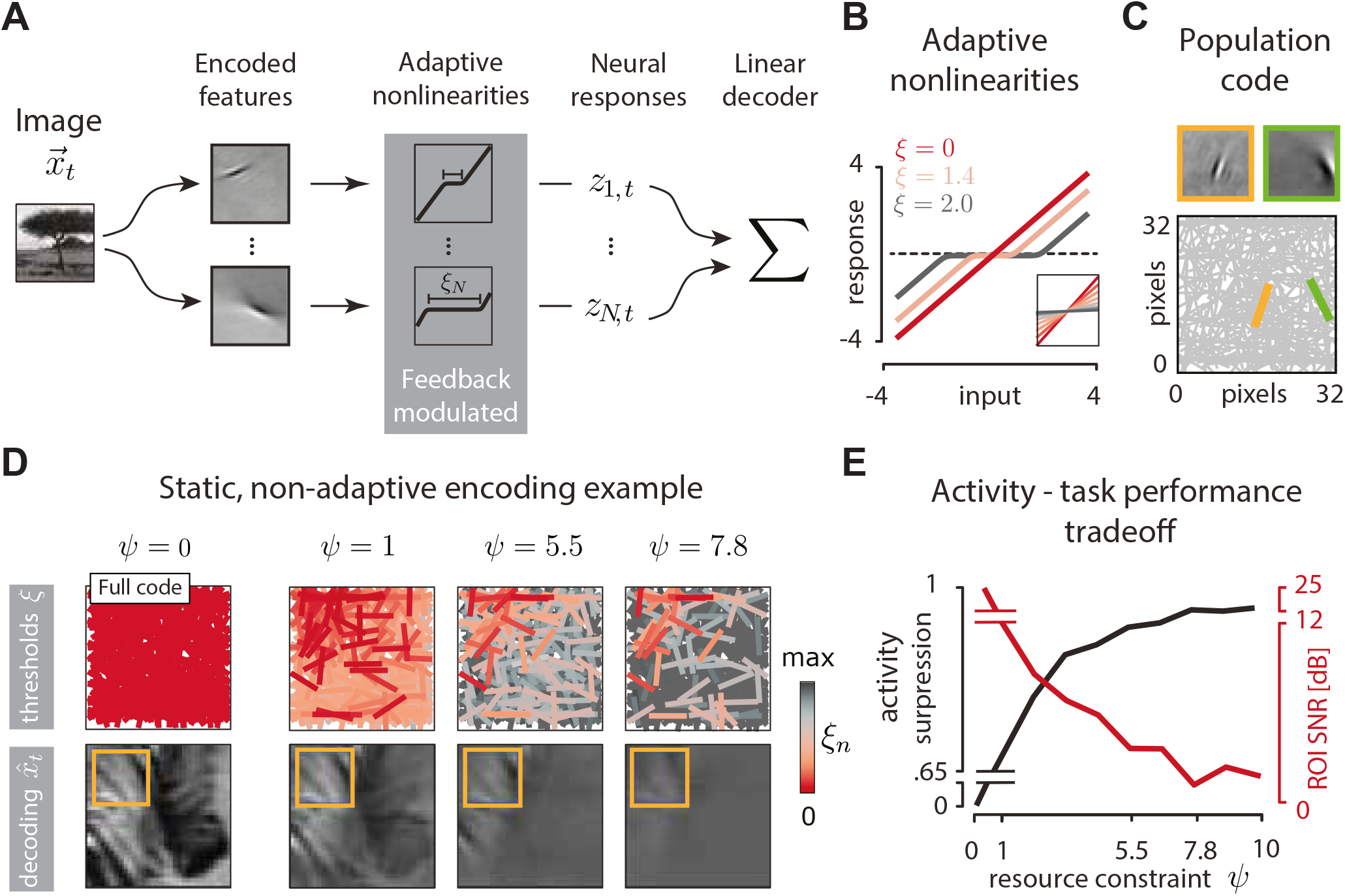
Adaptive population coding with nonlinearities. **(A)** An image 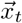 (32 × 32 pixel in size) is encoded by a population of *N* = 512 sparse coding model neurons, characterized by the represented features. Feature activations are transformed by adaptive nonlinearities with the threshold parameter *ξ*_*n,t*_. The resulting responses *z*_*n,t*_ are transmitted to the perceptual observer, which may use them to linearly decode the image and perform further task-specific computations. **(B)** Example adaptive nonlinearities for different values of the threshold parameter *ξ* (color). Inset: linear fits to nonlinearity outputs demonstrate that increasing the threshold *ξ* effectively decreases the neural response gain. **(C)** Visualization of the population code (bottom). The feature encoded by each model neuron is represented by a bar which matches that feature’s orientation and location. Two example features (top) are represented by bars of the corresponding color (bottom). **(D)** Left: an example image reconstructed using the standard sparse code (“full,” when all 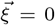). Orange frame marks a region of interest (ROI). Right, top row: three sensory populations optimized to reconstruct only the part of the image within the ROI, sorted by increasing attentional resource constraint *ψ*. Red intensity visualizes the value of the optimal thresholds *ξ*_*n*_ (red = low threshold and high gain; gray = high threshold and low gain). Right, bottom row: images linearly decoded from the corresponding sensory populations in the top row. **(E)** Activity of the neural population is increasingly suppressed (black line) and quality of ROI reconstruction (measured in dB SNR) decreases with increasing attentional resource constraint *ψ*.

The standard sparse coding model is capable of accurately reconstructing entire images, up to a single pixel, at minimal activity cost. Sparse coding can be viewed as an instantiation of efficient coding in a static, task-agnostic setup. We hypothesized that significant further efficiency gains would be possible if the sensory population could dynamically adjust its properties to encode only those image features required by the perceptual observer at any given moment.

We therefore extended the standard sparse coding model by transforming the output of each sparse feature with an adaptive nonlinearity (Fig. 2A). Each nonlinearity is controlled by a single parameter *ξ*_*n*_, which corresponds to an activation threshold (Fig. 2B). When *ξ*_*n*_ = 0, the response of the neuron *n* is equal to the activation predicted by the standard sparse coding. For *ξ*_*n*_ > 0 the neuron responds only when the activation exceeds a threshold determined by the value of *ξ*_*n*_. An increase of the threshold can be interpreted as a hyperpolarization of an individual neuron by feedback connections. Alternatively, an increase of the threshold can also be understood as an effective decrease in the neural gain (Fig. 2B, inset). This nonlinear transformation is reminiscent of smooth shrinkage, a well-known image denoising transform [33]. Neural nonlinearties can be dynamically modulated via feedback connections, as we describe more precisely below; what is essential here is that these nonlinearity adjustments allow the resulting neural responses *z_t,n_* to be sparsified beyond the standard, task-independent sparse coding. Mathematically, this is achieved by imposing an “attentional resource constraint” of strength *ψ* that penalizes high neural activity 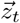 (see Eq (2), below). Finally, the neural responses are transferred downstream to the perceptual observer. Image decoding remains a simple, linear transformation.

To illustrate how this model population can selectively encode only the relevant features of a stimulus we consider a simple, static image encoding task (Fig. 2D). We optimize the nonlinearity parameters to reconstruct only a region of interest (ROI) of an image (Fig. 2D, orange frame). When the attentional resource constraint is inactive (*ψ* = 0), our model is equivalent to a sparse encoder, and the entire image can be reconstructed with high accuracy (Fig. 2D, leftmost column). For increasing values of attentional resource constraint *ψ*, the neuronal thresholds increase and “gain down” neurons that report on the image outside of the ROI (Fig. 2D, top row). While the quality of the overall image reconstruction deteriorates with increasing *ψ* (Fig. 2D, bottom row), the image within the ROI is preserved with high accuracy. The tradeoff between population activity suppression and ROI reconstruction accuracy as a function of the attentional resource constraint *ψ* is clearly visible (Fig. 2E).

This pedagogical example highlights how task-irrelevant features (here, image components outside of the ROI) can be suppressed in a sensory population to increase coding efficiency. Viewed from the image perspective, the sensory population is performing a lossy image compression, biased to maintain the image fidelity within the ROI. Here, the task is trivial, both because it deals directly with image reconstruction rather than inference, and because it is static. We now proceed to more complex and dynamic scenarios where the latent state of the environment and the stimulus statistics fluctuate. We expect that sensory neuron nonlinearities will have to dynamically adapt to the changing belief of the perceptual observer, as depicted in Fig. 1.

### Perceptual inference tasks

We consider three different probabilistic inference tasks that the perceptual observer carries out using the adaptive sensory code: object detection, target localization, and orientation estimation (Fig. 3A). These tasks correspond to traditionally defined types of attention: object-based attention, spatial attention, and feature-based attention, respectively.

**Figure 3:**
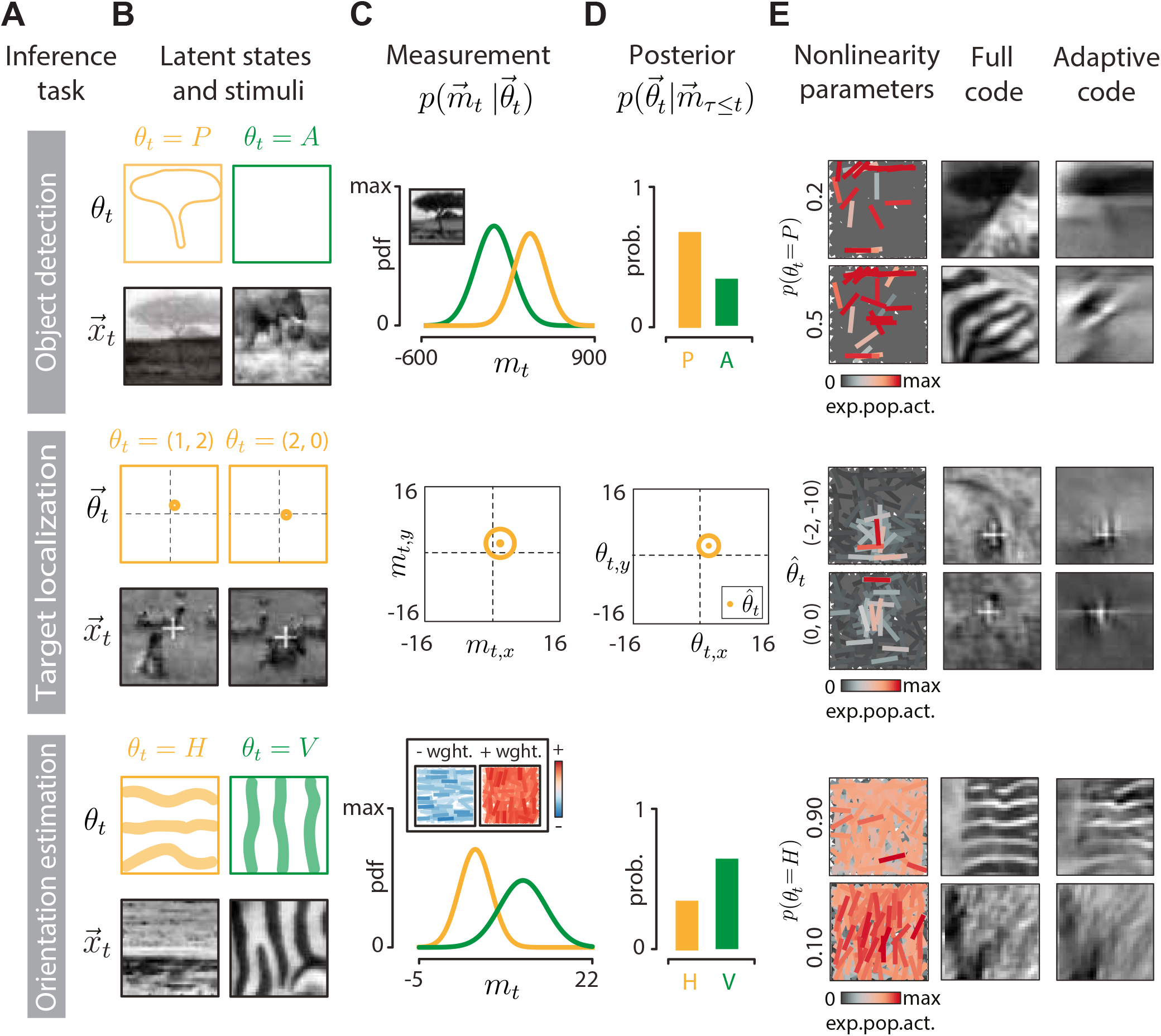
Perceptual inference tasks. **(A)** Rows correspond to individual inference tasks: object detection (top), target localization (middle), and orientation estimation (bottom). **(B)** Visualization of latent states 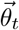 (top row of each panel, orange and green frames) and example stimuli 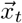 in each task (bottom rows of each panel, black frames). Top: tree present (orange) or absent (green). Middle: different white cross positions (orange dot). Bottom: orientation horizontal (orange) or vertical (green). **(C)** Measurements taken by the perceptual observer to infer the state of the environment. Top: a linear decoding of an image is projected onto a target “tree template” (inset) and noise is added. Measurements with object present (orange) and absent (green) follow different distributions. Middle: a linear decoding of an image is used to take a noisy measurement of the target position (orange dot = position estimate; orange circle = noise standard deviation). Bottom: logarithmically-transformed neural activity is projected onto a template (inset, blue and red = negatively and positively weighted neurons, respectively) and noise is added. Measurements of predominantly horizontal (orange) and vertical images (green) follow different distributions. **(D)** Posterior distributions. Top: probability of object being present (P, orange) or absent (A, green). Middle: probability of the visual target location (orange dot = MAP estimate; orange circle = covariance of the estimate). Bottom: probability of the image being predominantly horizontally (H, orange) or vertically (V, green) oriented.**(E)** Top row, left column: population activity for two different observer belief levels that the tree is present. Top row, middle column: two images decoded using the full code optimized for image reconstruction. Top row, right column: two images decoded using the adaptive code with the activity shown in the left column. Middle and bottom rows: analogous to the top row, but for target localization and orientation estimation, respectively. Throughout, the neural population is visualized using the expected neural activation (colorbar; see Methods).

For each task, the perceptual observer performs a sequence of computations outlined in Fig. 1 at each time step. First, the observer uses a representation of the stimulus in the form of population activity vector 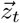 to perform a “measurement” 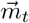 of the stimulus feature required to infer the latent variable of interest. The measurement consists of evaluating a task-dependent function *f* over the population activity vector, i.e., 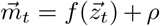, where *ρ* is additive Gaussian noise. Second, the measurement 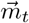 is used in a Bayesian update step to compute the distribution over the latent state of the environment 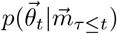, and the predictive distribution of future stimuli 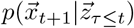. Third, the predictive distribution is used to select optimal values for the neural nonlinearities, to be conveyed to the sensory population via top-down feedback (see Methods for details).

#### Object detection

The goal of the object detection task is to infer whether a specific object is embedded in the current image or not (Fig. 3A, B, top row). The latent state of the environment follows a random correlated process to switch between “object present” (*θ* = *P*) and “object absent” (*θ* = *A*). The observer linearly decodes the image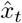 and computes the measurement *m_t_* by projecting the decoded image onto the object template. The measurement *m*_*t*_ follows a different distribution, depending on whether the object is present or absent in the scene (Fig. 3C, top row). The posterior distribution is characterized by a single number, the probability of object present *p*(*θ* = *P*)(Fig. 3D, top row).

#### Target localization

The goal of the target localization task is to infer the position of a moving visual target – a white cross – embedded in the background of a natural movie (Fig. 3A, B, middle row). The observer linearly decodes the image to extract a noisy measurement of the position of the target, by computing cross-correlation with the target template (Fig. 3C, middle row; see Methods). This noisy measurement, combined with observer’s knowledge of the target dynamics, is used to estimate the current position of the target along the two spatial coordinates 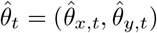 (Fig. 3D, middle row).

#### Orientation estimation

The goal of the orientation estimation task is to determine whether the current stimulus is predominantly horizontally or vertically oriented (Fig. 3A, B, bottom row). These two classes of images were first discovered via unsupervised learning ( see Methods). The latent state of the environment follows a random correlated process to switch between “horizontal” (*θ* = *H*) and “vertical” (*θ* = *V*). The observer projects the magnitudes of neural responses 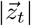 onto a discriminative template, without decoding the image first, to obtain the measurement *m*_*t*_ (Fig. 3C, bottom row; see Methods for details). The measurement follows different distributions for horizontally and vertically oriented images (Fig. 3C, bottom row). The posterior distribution is characterized by a single number, the probability that the environment is in the horizontal state *p*(*θ* = *H*) (Fig. 3D, bottom row).

### Top-down feedback for adaptive coding

To complete our model, we must close the loop and specify the procedure by which the perceptual observer adapts nonlinearity thresholds in the sensory population via top-down feedback. Following the normative approach, we assume that such adaptations are mathematically optimal and proceed to work out the predicted consequences. The question of how realistic neural circuits could implement or approximate the required optimality computations is clearly important but beyond the scope of present work.

We start by assuming that at each time step a predictive distribution over future stimuli,

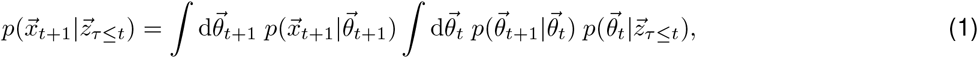

can be computed to derive the optimal nonlinearity parameters, 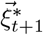, which, in turn, are used to encode the future stimulus. These optimal parameters 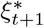 are chosen at every time step to minimize the following cost function:

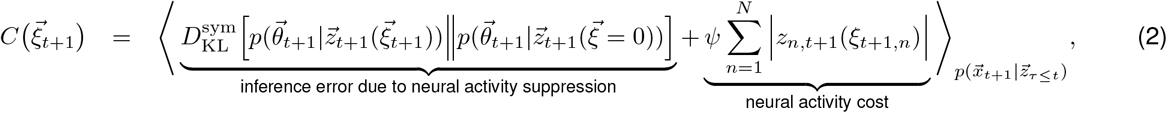

where 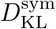 is the symmetrized Kullback-Leibler divergence. We relied on symmetrized variant of the KL divergence because of its conceptual similarity to other error measures such as reconstruction error. The essence of the framework outlined here is however not dependent on this precise choice.

The cost function in Eq (2) is at the core of our approach. The first term corresponds to the error in inference induced by image compression due to suppression of the neural activity via adaptive thresholds (see Methods): this term is small in expectation when the task-relevant predictive information can be retained (at low threshold values). The second term is the neural activity cost, where *ψ* is the attentional resource constraint: this term is small when the predicted neural activations will be sparse (at high threshold values). By minimizing the cost function *C*, the system balances the two opposing objectives and minimizes the error in latent state inference while reducing the amount of neural activity beyond the limit set by standard sparse coding (*ψ* = 0).

Optimal thresholds depend on the current task, the strength of the attentional resource constraint *ψ* and, crucially, on the time-changing perceptual belief of the observer. The dynamics of this belief induce temporal changes in the structure of the sensory code. This interplay is illustrated in Fig. 3E. In the object detection task (Fig. 3E, top panel) only the neurons which encode the silhouette of the object are modulated, while the rest of the population remains suppressed to minimize activity. When the observer does not believe that the tree is present in the scene (i.e., *p*(*θ* = *P*) is low; Fig. 3E, top panel, top row) only a minimal set of neurons remains active, in order to encode the outline of the tree should it suddenly appear. This is evident when comparing the image decoded from the full code with that from the adaptive code: in the latter case, only the shape of the tree is retained while the rest of the image detail is compressed out. When the uncertainty about the presence of the object increases (i.e., *p*(*θ* = *P*) = 0.5), the sensory population must preserve additional image features to support the perceptual task (Fig. 3E, top panel, bottom row).

Similar reasoning applies to the orientation estimation task (Fig. 3E, bottom panel), where only the neurons encoding the relevant image orientations remain active and modulated by the observer. While the images reconstructed from the adaptive code lose a lot of spatial detail, they retain the global “gist” which enables the observer to identify their dominant orientation.

The influence of perceptual belief on the sensory encoding is perhaps most clearly apparent in the target localization task (Fig. 3E, middle panel). Here, the sensory population encodes only that region of the image where the perceptual observer believes the target is expected to move in the next time step. This task can be seen as a dynamic generalization of the ROI encoding example of Fig. 2D. As the target moves, the observer extrapolates this motion into the future and encodes information just sufficient to confirm or rectify its prediction, while suppressing the rest of the image. This results in an attentional phenomenon that closely resembles a moving spatial “spotlight” of high visual acuity.

### Adaptive coding enables accurate inference with minimal neural activity

How do adaptive codes navigate the tradeoff between minimizing neural activity and maximizing task performance? We simulated perceptual inference in dynamic environments over multiple time steps for all three tasks (Fig. 4A). Adaptive coding results in drastic decreases of neural activity in the sensory population compared to the standard sparse coding (Fig. 4B). Adaptive coding furthermore reveals interesting task-specific dynamics of population activity, locked to the switches in the environmental state. For example, in the object detection and orientation estimation tasks (Fig. 4B, top and bottom panels, respectively) the neural activity is significantly decreased in “absent” and “horizontal” environmental states, respectively. This is because the sensory system needs to extract different kind of information to support downstream inferences in different environmental states. In contrast, the standard sparse code maintains a roughly constant level of activity (Fig. 4B, red lines).

**Figure 4:**
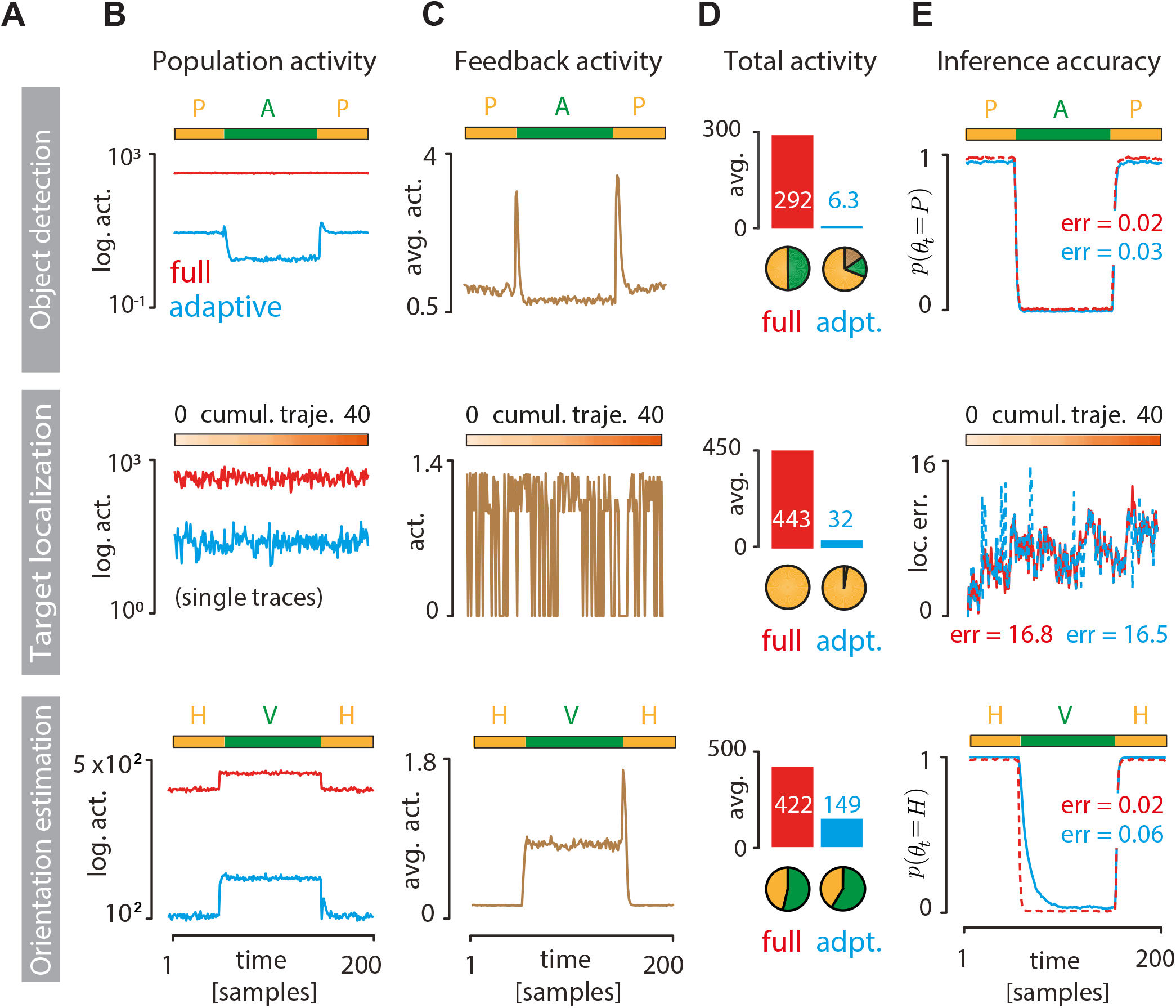
Adaptive coding significantly reduces activity cost with minimal impact on inference accuracy. **(A)** Rows correspond to inference tasks: object detection (top), target localization (middle), and orientation estimation (bottom). **(B)** Sensory population activity ⟨|*z*_*n,t*_|⟩*_n_* in the standard sparse code optimized for image reconstruction (red = full code) or for a particular task (blue = adaptive code). Activities in object detection (top) and orientation estimation (bottom) tasks were averaged over 500 switches between different states of the environment. For the target localization task (middle), we plot an short non-averaged activity segment (200 time steps out of a 10^4^ time-step simulation; see Methods).**(C)** Same as B but for feedback activity required to adapt the nonlinearities in the sensory population (see Methods). **(D)** Time-averaged activity of the full code (red bars) and adaptive code (blue bars). Pie charts show the total activity decomposed into contributions from two different environmental states (green and orange; top and bottom row only) and feedback (brown; adaptive codes only). An important signature of adaptive coding is the asymmetric distribution of activity across environmental states. **(E)** Inference accuracy (red = full code; blue = adaptive code). Estimates of the environmental state (“object present” in object detection task, top; “orientation horizontal” in orientation estimation task, bottom) were averaged over 100 environmental switches. For the target localization task (middle), inference accuracy is measured as mean squared error between the true and inferred position of the target cross. Text insets display the average inference error in each task (see Methods).

We also quantified the cost of top-down feedback signaling (Fig. 4C). In our model, feedback activity is commensurate with the amplitude and frequency of posterior belief updates in the perceptual observer (see Methods), making feedback activity patterns strongly task-specific. In the object detection task, feedback activity peaks briefly during switches between environmental states (Fig. 4C, top panel). In the orientation estimation task, the belief of the perceptual observer fluctuates strongly when vertical orientation dominates, leading to elevated feedback activity (Fig. 4C, bottom panel). Since the signal statistics are more homogeneous in the target localization task, feedback activity (when non-zero) stays within a tight interval (Fig. 4C, middle panel).

Despite the additional cost of feedback signaling, the total activity of adaptive codes is drastically lower compared to the full sparse code, sometimes by more than an order of magnitude (Fig. 4D). This dramatic reduction does not significantly impact the accuracy of the inferences (Fig. 4E). Average trajectories of the posterior probability for the object detection and orientation estimation tasks are very similar (Fig. 4E, top and bottom panels). In the target localization task, the instantaneous error of the target location estimate using the adaptive code closely follows the error of the full code (Fig. 4E, middle panel). For all tasks, the time-averaged error values are comparable between the adaptive and the full code. Taken together, this demonstrates that adaptive coding enables accurate inferences while dramatically minimizing the cost of neural activity in the sensory population.

### Statistical signatures of adaptive coding

Dynamic adaptation significantly changes the statistical structure of a sensory code. The most prominent change is a large increase in the sparsity of the adaptive code compared to the standard sparse code across all tasks (Fig. 5A,B). This finding is consistent with the observed suppression of average neural activity (Fig. 4D). These two phenomena are, however, not exactly equivalent. Sparsity of neural responses (as measured by kurtosis) can be increased in many ways [26], and each would result in suppression of the average activity. In our case, sparsity increase in the adaptive code is induced specifically by a complete suppression of a subpopulation of neurons, resulting in the high spike at zero in the neural response distribution (Fig. 5A).

**Figure 5:**
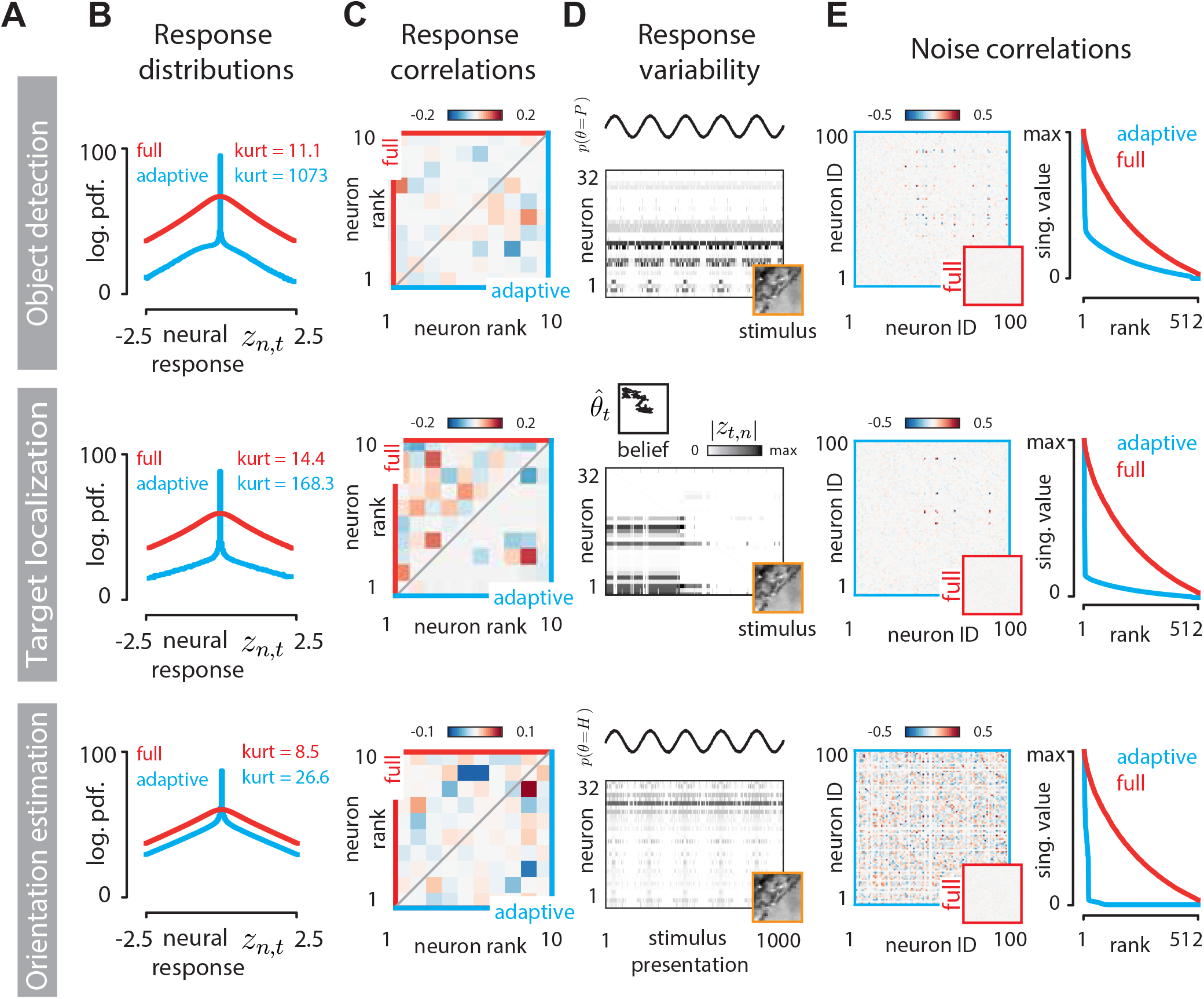
Statistical differences between the adaptive code and the standard sparse code. **(A)** Rows correspond to inference tasks: object detection (top), target localization (middle), and orientation estimation (bottom). **(B)** Distributions of neural responses *z*_*t,n*_ for the standard sparse code code optimized for image reconstruction (full, red) and the adaptive code (blue); kurtosis as a measure of sparsness is displayed in inset. **(C)** Pairwise correlations of 10 example neurons whose activity is modulated by the task (different for each task). Correlations were computed over the entire stimulus trajectory used to generate plots in Fig. 4. Upper triangle (red) of correlation matrices corresponds to the full code, bottom triangle (blue) to the adaptive code. **(D)** Belief-induced response variability in the adaptive code. Neural activation (grayscale proportional to |*z*_*n,t*_|^0.5^) for 32 example neurons chosen separately for each task, exposed to 1000 presentations of the same stimulus (orange frame). Response variability at fixed stimulus originates from the fluctuations in the internal belief of the perceptual observer (top part of each panel). Here, these fluctuations are simulated as sinusoidal variations in the probability of environmental state (object detection and orientation estimation tasks; top and bottom row, respectively), or a random walk trajectory of the target for the localization task (middle row). **(E)** Belief-induced noise correlations in the adaptive code. Left column: correlation matrices of the same 100 neurons computed from responses to stimulus presentations displayed in D. Right column: scaled singular values of correlation matrices of the adaptive code (blue). We compared this spectrum to the standard sparse coding in which a small amount of independent Gaussian noise is added to each neural activation. In this case, noise correlation singular values scale with the noise magnitude (not shown), and their normalized singular spectrum is denser (red) compared to that of the adaptive code.

Coordinated top-down modulation of individual neurons leaves its imprint also on the collective statistics of the population activity. For example, different perceptual tasks engage different neurons and, among them, induce different patterns of pairwise correlation. This effect becomes apparent when we focus on a subset of neurons active in a task and compare their correlated activity under standard sparse code or under the adaptive code. In the standard sparse code, neural correlations are inherited solely from the stimulus (Fig. 5C, top submatrices, red frame). In an adaptive code, they are additionally modulated by the task, leading to a very different correlation pattern (Fig. 5C, bottom submatrices, blue frame).

Changes in the stimulus are not the only factor which drives response variability in the visual cortex. Cortical responses are notoriously unreliable and can fluctuate widely over multiple presentations of the same stimulus [34], giving rise to “noise correlations” among sensory neurons [35]. Patterns of noise correlations can be task-specific and driven by feedback [3]. Our framework provides a new normative hypothesis about the origin and functional relevance of response variability and noise correlations. In our model, neurons generate different responses even at fixed stimulus when the neural nonlinearities change due to fluctuations in the internal state of the perceptual observer. For example, at the beginning of each target localization trial – even though the stimulus is the same – the perceptual observer may have a different prior belief about where the target is, possibly influenced by preceding history of the neural dynamics or sampling noise that leads to stochastic information accumulation about target position. Trial-to-trial differences in this internal belief will result in a variable allocation of resources in the sensory population as directed by the perceptual observer via top-down feedback, leading to strong noise correlations.

We simulated such a scenario by exposing our model to multiple presentations of a single stimulus, identical across the three tasks, while enabling the perceptual belief to vary. A clear pattern of response variability to multiple presentations of the same stimulus is visible in each case (Fig. 5D). This task-specific and feedback-driven response variability manifests in distinct noise correlation structures (Fig. 5E, left column). For the adaptive code, the noise correlation matrix is dominated by a small number of modes, reflecting a low-dimensional fluctuating internal state of the perceptual observer. In contrast, noise correlations are expected to be exactly zero for the standard sparse code. If independent noise is purposefully introduced into the standard sparse coding units (see Methods), the singular value spectrum is much denser than for the adaptive code (Fig. 5E, right column), indicating that the presence low-rank noise correlations differentiates between adaptive and full sparse codes. This observation is consistent with the experimentally observed low-dimensionality of task-specific correlations in the visual cortex [3].

Taken together, adaptive code is predicted to feature: first, a sparser response distribution compared to the standard sparse code; second, task-dependent response correlations compared to task-independent correlations for the standard sparse code; third, prominent yet low-rank noise correlations compared to zero noise correlations for the standard sparse code.

### Adaptive coding reproduces attentional phenomena in the visual cortex

To check whether our normative approach accounts for salient attentional phenomena observed experimentally, we qualitatively compared the properties of the adaptive coding model with the signatures of top-down attention measured in the visual cortex (Fig. 6). For this comparison we used the target localization task due to its similarity to the established spatial attention paradigm [1]. While our model is intended primarily as an idealization of the primary visual cortex, below we compare it to data on target localization from both V1 and V4; we believe this is justified since the observed effects have been observed across the visual hierarchy from V1 to V4 and IT, and since these effects are a consequence of generic computational principles that are likely shared across cortical regions.

**Figure 6:**
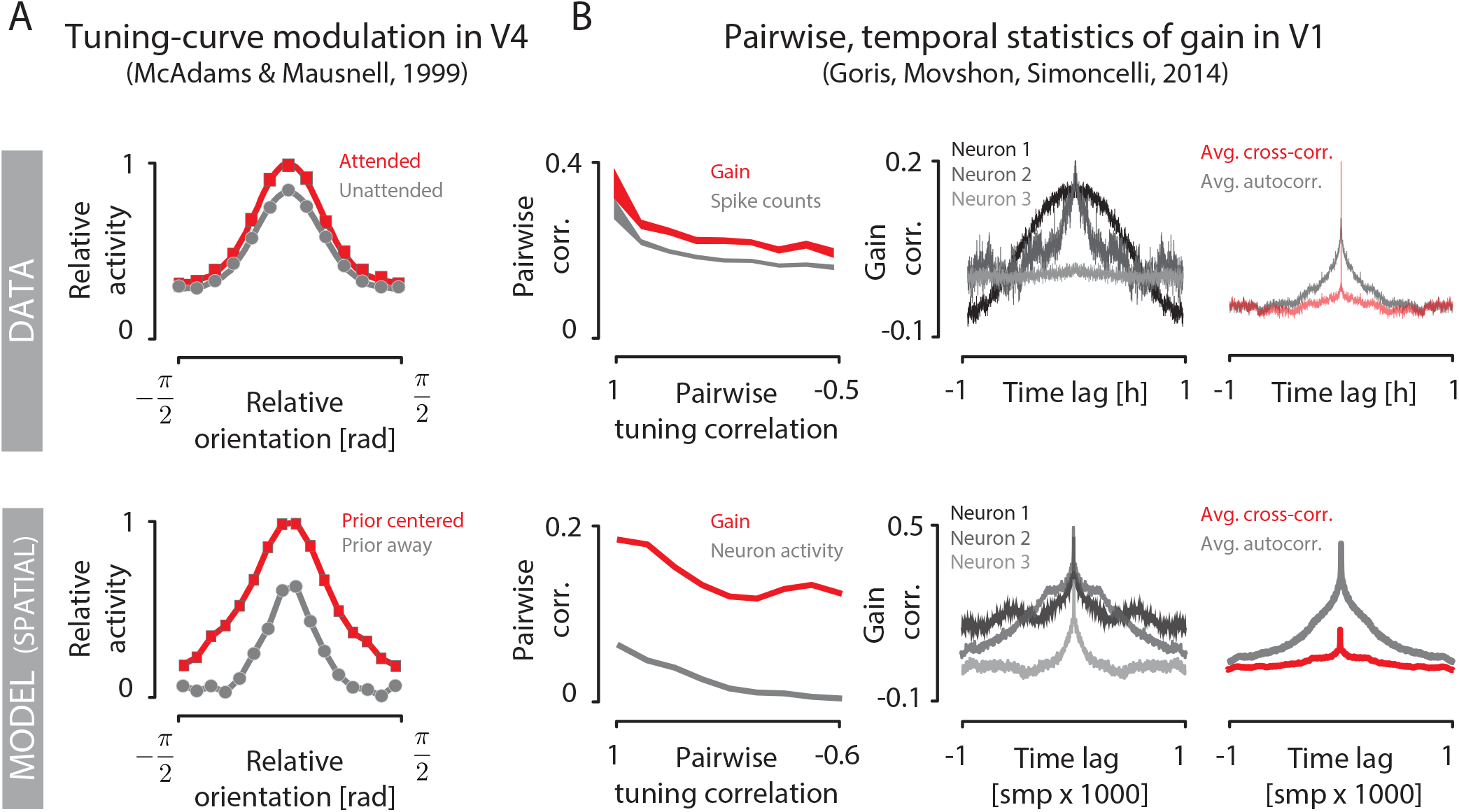
Comparison of adaptive coding model for target localization to experimental data. **(A)** Population tuning curves in macaque V4 in an attended (red) and unattended (gray) conditions (top panel, replotted from [14]). Model prediction (see main text) reproduces the modulation of tuning curves (bottom panel). **(B)** Pairwise correlation of internal gain signals (red) and neural activity (gray) as a function of tuning correlation in macaque V1 (top left) is reproduced by the model (bottom left; see main text). Measured gain autocorrelation functions for three example neurons (top middle) resemble optimal gain dynamics in the model (bottom middle). Average gain autocorrelation function (gray) and average pairwise gain cross-correlation function (red) are reproduced by the model (data figures -courtesy of Robbe Goris [19] top right; model bottom right).

A prominent hallmark of spatial attention in the visual cortex is the modulation of population tuning curves [14]. Orientation-selective neurons whose receptive fields are located in the attended part of the scene respond more strongly to preferred stimuli than neurons encoding unattended parts of the scene (Fig. 6A, top). Our model optimized for the target localization task reproduces this phenomenon (Fig. 6A, bottom). When the perceptual observer expects the target to be present at a particular image location, it increases the gain of neurons reporting on that location. We interpret this as equivalent to top-down attention being directed towards that location, which allows us to extract from our model a “prior-centered” tuning curve comparable to the “attended” experimental condition. This is to be compared with the “baseline” tuning curve comparable to the “unattended” experimental condition, computed using neural gain averaged over long periods of time (see Methods). We note that this spotlight-like gain modulation was not engineered in any way into our model; instead, it emerged from a generic principle that optimizes perceptual inference under coding cost constraints.

Another prominent signature of attentional processing in the visual cortex concerns neural response variability, which can be conveniently separated into sensory drive and gain dynamics [19, 20]. Specifically, gain dynamics might reveal internal brain states related to arousal and attention [19]. Here we compare the statistical structure of gain dynamics predicted by our normative model with the measurements in the visual cortex (Fig. 6B). Because changes in effective neural gain are linked to changes in activation thresholds *ξ* in our setup (Fig. 2B), we focus on predicted neuron-to-neuron correlations in threshold dynamics as well as individual neuron threshold autocorrelation function (see Methods). Clear similarities emerge. First, observed correlations of gain and neural activity decay with decreasing correlation of neuronal tuning, as predicted by our model; furthermore, the activity correlation is consistently lower than the gain correlation, also as predicted (Fig. 6B, left column). Second, a broad spectrum of temporal dynamics for the gain of individual neurons is observed in the sensory population: from long temporal correlations to almost instantaneous decay, which is correctly reproduced by our model (Fig. 6B, middle column).When averaged over multiple neurons, the gain autocorrelation function shows a smoothly decaying profile. In contrast, the average cross-correlation in gain across pairs of neurons reveals no preferred temporal relationship and decays essentially instantaneously, which is correctly reproduced by our model (Fig. 6B, third column). This finding is surprising, because – at least in the model – global coordination by top-down feedback could be expected to lead to non-zero temporal correlations in gain.

### New predictions of adaptive coding

Previous theoretical work established a link between perceptual uncertainty about the state of the environment and the influence of stimuli on the perceptual belief [31]. In brief, when a Bayesian perceptual observer is highly certain about the value of a latent state of the environment (strong prior), subsequent sensory signals will only have a small influence over its belief (the posterior will be similar to the prior). In contrast, when the observer is highly uncertain, any individual stimulus can sway the observer’s belief by a large margin (the posterior can differ significantly from the prior). This reasoning leads us to the following hypothesis: efficient sensory systems gain down stimulus encoding in states of high perceptual certainty and gain up encoding in states of high perceptual uncertainty.

We tested this hypothesis in our model. Across all tasks, increases in perceptual uncertainty lead to increased population activity (Fig. 7A,B). In contrast, standard sparse coding is not modulated by uncertainty and maintains its activity at a high baseline required to reconstruct the stimuli in full.

**Figure 7:**
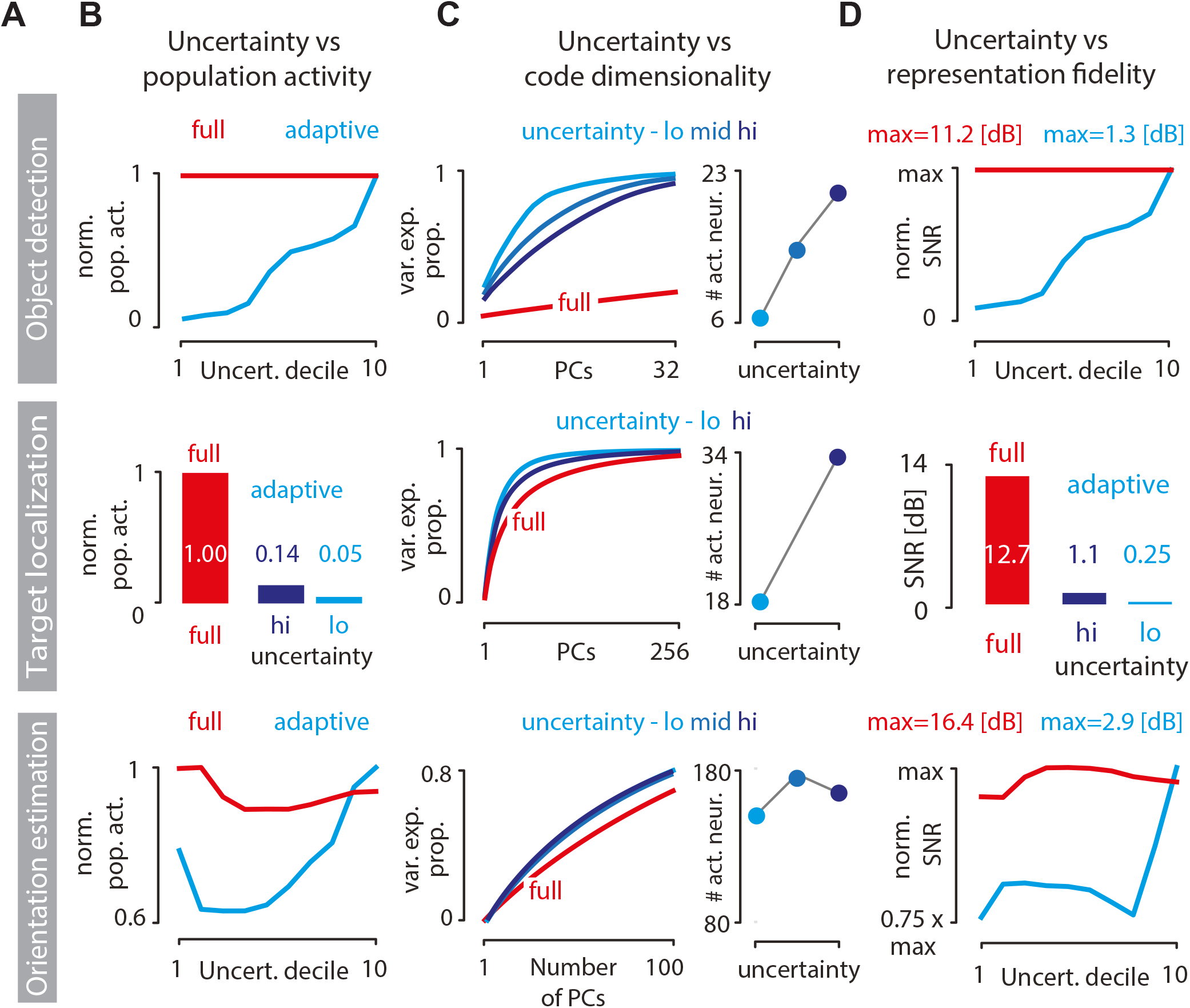
Predicted changes in the adaptive code when perceptual uncertainty is manipulated. **(A)** Rows correspond to inference tasks: object detection (top), target localization (middle), and orientation estimation (bottom). **(B)** Normalized population activity as a function of perceptual uncertainty for the standard sparse code (red = full code) and the adaptive code (blue). Uncertainty in object detection (top) and orientation estimation (bottom) tasks was binned into deciles (see Methods). Uncertainty in the target localization task (middle) is plotted for two levels of measurement noise (dark blue = high noise; light blue = low noise). **(C)** Dimensionality of the adaptive code can increase with increasing perceptual uncertainty (left column). Shown is the proportion of variance in total neural activity explained as a function of the number of principal components (red = full code; light blue = adaptive code at low uncertainty; medium blue = adaptive code at intermediate uncertainty; dark blue = adaptive code at high uncertainty; see Methods). Increase in code dimensionality is correlated with the number of active neurons at different levels of uncertainty (right column). **(D)** Same as B but showing the normalized SNR of the image reconstruction at different perceptual uncertainty levels.

Does perceptual uncertainty affect only the total amount of neural activity or also its statistical structure? To answer this question, we assessed the dimensionality of sensory population activity with principal component analysis, and analyzed it as a function of the entropy of the prior that the perceptual observer holds about the environmental state (see Methods). In two out of three perceptual tasks, progressively uncertain observer engages increasing numbers of neurons (Fig. 7C, right column top and middle panels), which affects the dimensionality of the sensory code. When the observer is highly certain, few principal components suffice to explain the population activity; as perceptual uncertainty grows and progressively more neurons are engaged via top-down feedback, the dimensionality of the code increases, but always remains bounded by the dimensionality of the full sparse code (Fig. 7C). These changes are mirrored in the fidelity of stimulus reconstruction that can be read out from the sensory population (Fig. 7D): as perceptual uncertainty grows, incoming stimuli are increasingly relevant for inference and more sensory resources are deployed to encode the stimuli, leading to improvements in stimulus reconstruction.

These results generate two new experimental predictions. First, the average firing rates and the dimensionality of neural activity in the visual cortex should increase during periods of high perceptual uncertainty about the state of the environment. This could be tested, for example, in the target localization paradigm, by comparing experimental conditions in which the target object follows a more vs less predictable trajectory, or where the target is embedded at a higher vs lower contrast in a structured background. To control for sensory confounds and isolate specific effects of perceptual uncertainty, it should be possible to design stimulus protocols where the perceptual task is always performed with an identical probe stimulus, but where perceptual uncertainty was manipulated by prior exposure to different priming stimuli. These predictions echo recent findings that link neural gain variability to perceptual uncertainty induced by manipulations of low-level image statistics [36]. This link between uncertainty and variability is also qualitatively captured by our model (see Supplemental Fig. S2).

Second, our results predict that disruption of top-down signaling should decrease the variability of responses in the sensory population. According to our model, the frequency and strength of top-down feedback signaling grows with perceptual uncertainty and the frequency of perceptual belief changes. As a consequence, it should be possible to compare the activity of the intact sensory population with the activity of the sensory population where top-down feedback was interrupted via mechanical, pharmacological or optogenetic means, under stimulus or task conditions that induce large fluctuations in perceptual uncertainty. Disrupted feedback should decrease variability in the sensory population and stabilize its statistics.

## Discussion

Attention has long been a subject of theoretical research and numerous theories of its origin and functional relevance have been proposed [23, 24, 37–39]. In this work we suggest that several open questions about attention—about its phenomenology, its effects on the neural code, and its functional origins, as laid down in the Introduction—are interrelated and fall within the purview of a single conceptual framework that synthesizes two canonical theories of neural computation: optimal perceptual inference and efficient coding [31, 40, 41].

To make these ideas concrete, we develop a model of sensory coding in the visual cortex that is applicable to dynamic and possibly non-stationary scenarios. We demonstrate that attention-like phenomena emerge as a consequence of moment-to-moment adaptations in a resource-limited sensory code optimized to efficiently learn about the states of the environment. Such “optimal adaptive coding” reproduces a number of observations previously attributed to attention: emergence of spatial spotlight, tuning curve modulation, gain dynamics, task-dependence of neural correlations, and response variability manifesting as noise correlations. We furthermore suggest that different kinds of attention should not be thought of in terms of distinct neural mechanisms, but rather as a natural consequence of universal computational principles.

Our framework also bears on a puzzling paradox at the heart of how we understand sensory systems. On the one hand, perception and attention seem to rely on coarse, high-level properties of visual scenes which are encoded selectively depending on the goals and internal states of the brain [42, 43]. On the other hand, neurons in the sensory periphery encode signals at the physical limits of precision, right up to individual photons [44]. Why invest in such precision if the information is subsequently not used to guide perception or behavior? Our model shows that adaptive sensory systems which possess the ability to accurately encode the entire image with a single pixel accuracy can also dynamically partition this sensory information, into the task-relevant part to be extracted and the task-irrelevant part to be suppressed. Precise sensory representations can thus be maintained at a higher cost only when needed; when they suffice for the task, coarse sensory representations are preferred for their efficiency.

### Relationship to other theoretical frameworks

The hypothesized role of top-down feedback in our approach differs from previous proposals. In hierarchical predictive coding, feedback conveys predictions of the higher-order stimulus structure to the sensory population, so that sensory neurons only need to encode the difference between such predictions and the true stimuli [45]. This establishes a complete and efficient representation of the signal. Top-down feedback has a related role in hierarchical Bayesian inference [46], where it conveys information about higher-order statistical regularities of the stimulus to “explain away” lower-level activity. Sensory hierarchy instantiates a complete representation of the stimulus, with higher levels corresponding to progressively more abstract features of the signal [27]. Importantly, in both of these classical approaches the system needs to perform multiple feed-forward and feedback passes to establish the complete stimulus representation.

In contrast to the above theories that retain stimulus detail up to the pixel level, our model of adaptive coding involves a (potentially lossy) compression of sensory stimuli. Here, top-down feedback does not provide the values needed for prediction subtraction (in predictive coding) or for explaining away (in hierarchical Bayesian models); in fact, top-down feedback conveys no stimulus information, at least not directly. Instead, feedback conveys the optimal “system settings” for the lossy encoder (e.g., nonlinearity parameters for the sensory population), based on predictions of the perceptual observer. In this setting, the sensory system does not require multiple feed-forward and feedback passes to establish the stimulus representation. As a consequence, neural resources devoted to coding and time devoted to transmission of sensory information are dramatically reduced. This efficiency comes at a cost – the resulting representation is less robust and unexpected environmental changes may lead to dramatic errors in perceptual inference. Taken together, the adaptive coding regime instantiated by our model offers a perspective on the role of top-down feedback in sensory systems that is complementary to previous work.

In our approach attention-like processing emerges as a consequence of optimizing a general-purpose objective function. The model derived here fits in a broader tradition of deriving sensory codes from principles of efficient coding [26, 47]. Phenomena such as the spatial spotlight or enhancement of vertical orientations are therefore a “side-effect” of this optimization, rather than a goal in itself. The objective function considered here is reminiscent of other normative objectives for sensory coding in dynamic environments, such as maximization of predictive information [48]. There, the goal of the sensory system is to extract information predictive of future values of the stimulus. We consider the case where encoded information is expected to be relevant for a latent-variable inference and not the raw values of the stimulus, suggesting further connections to the information bottleneck framework [29, 49]. We foresee exploring the relationship between these objectives in future work.

Our model provides an intriguing link between attentional mechanisms and a wide body of work on the origins and effects of neural correlations. It has been suggested that neural correlations might be a consequence of shared noise sources and circuit connectivity [50], or that they may reflect hierarchical perceptual inference [51] which can be instantiated via probabilistic sampling [51–53]. We do not see our model as being mutually exclusive with these proposals: adaptive coding could contribute partially, along with other mechanisms, to the task-specific changes in experimentally measured neural correlations. An important debate concerns the impact of neural correlations on information transmission in sensory systems [54, 55]. Our framework provides a nuanced perspective on this issue. Here, changing correlation patterns reflect the compression of the stimulus, i.e., the reduction of the total encoded stimulus information. Removed stimulus information is however unnecessary from the perspective of the task to be solved and the momentary perceptual belief.

### Caveats and future work

Our work crucially depends on the observer using the correct statistical model of the environment and its dynamics. Dramatic reduction of neural activity cost with a negligible impact on inference quality cannot be achieved by a “mismatched” observer, which uses an incorrect model, operates under incorrect assumptions, or fails to correctly compute optimal thresholds. To perform accurate inferences, a mismatched observer should encode stimuli in higher detail and should therefore decrease the attentional resource constraint *ψ*. We foresee the possibility of dynamically controlling this parameter: attentional resource constraint could start at a low value which would progressively increase as the observer converges to the correct model of the environment. The optimal dynamics of the attentional constraint itself (“meta-attention”) is a subject of future work.

Our model makes a number of idealizations about the sensory neuron population. Firstly, we assume that adaptive nonlinearities are applied to the output of the sparse coding population, where lateral inhibition plays a crucial role in forming the code [26, 32]. In this scenario, activations *s*_*n,t*_ of the sparse-coding algorithm correspond to subthreshold potentials [56]. Neural firing is computed in a separate step, by transforming these potentials with a thresholding nonlinearity. We envision other possible mechanisms where suppression of unnecessary neural activities occurs simultaneously with the computation of the sparse code, for example, by manipulating sparsity constraints of individual neurons. Secondly, our neural activity is real-valued, making direct quantitative comparisons with real data impossible for features such as response variability; this issue could be addressed by extending the model with Poisson spike generation. Lastly, we assume instantaneous top-down feedback, whereas real neural circuits may suffer from transmission delays that could detrimentally affect the code performance.

Despite the assumptions described above, our key insights should not depend on detailed modeling choices. Compression of sensory signals could be achieved with different types of nonlinearities, or transformations such as divisive normalization and multiplicative scaling [57, 58]. Similarly, stimulus could be represented by alternative schemes, e.g., by neural sampling [56]. Inference carried out by the perceptual observer also need not be explicitly probabilistic [59]. The only essential component of our model is the feedback loop that dynamically adapts the sensory code to the demands of the perceptual observer. This provides the necessary theoretical link between the dynamics of attentional processing, efficient coding, and perceptual inference.

## Methods

### Adaptive coding model of natural images

#### Spare coding model of V1

Standard sparse coding model [26] represents image patches *x*_*t*_ with a population of *N* neurons, each of which encodes the presence of a feature 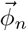 in the image. Given activations of individual neurons *s*_*n,t*_, the image patch can be linearly decoded as:

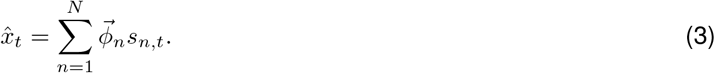

Basis functions *ϕ* are optimized to jointly minimize the reconstruction error and the cost of neural activity (or, conversely, to maximize sparsity):

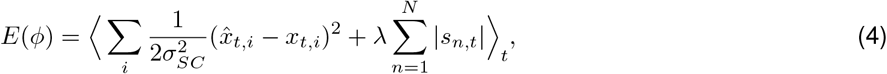

where *λ* is the sparsity constraint, 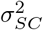 is the noise level, *i* indexes image pixels, and *t* indexes individual images in the training dataset. We optimized a set of *N* = 512 basis functions using the standard SparseNet algorithm [26] which iteratively alternates between minimizing Eq. (4) with respect to basis functions *ϕ* and coefficients *s*. During learning, we fix ||*ϕ*_*n*_||^2^ = 1 for every *n*.To learn neural receptive fields we used a dataset of 5·10^4^ 32 × 32 pixel image patches (standardized to zero mean and unit variance for each patch) randomly drawn from natural movies of the African savannah [60], which were reduced to 512 dimensions using Principal Component Analysis. We learned the sparse features *ϕ* using *λ* = 1 and 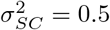 we then fixed features *ϕ* for all subsequent analyses.

#### Adaptive nonlinearities

We extended the sparse coding model by applying pointwise nonlinearities to sparse coding outputs. After encoding an image patch 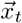 we transformed the activations of individual neurons *s*_*n,t*_ into responses *z*_*n,t*_:

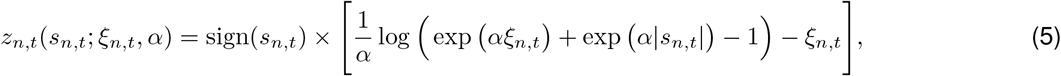

where *ξ*_*n,t*_ is the threshold value and *α* = 10 is a constant parameter. This nonlinearity is a smooth and differentiable shrinkage operator proposed in Ref [61]. Thresholds *ξ*_*n,t*_ are individually set for each neuron at each time point to encode only these features of the image which are required to perform the perceptual inference.

#### Visualization of nonlinearity parameters

To compare different threshold settings *ξ* in the sensory population across tasks, perceptual beliefs and stimulus distributions, we visualized the expected neural activity of neuron *n* at time 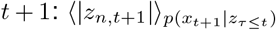. This quantity, which we typically display in color code, would correspond to experimentally observable expected activity of neuron *n*.

#### Cost of feedback activity

We assume that the feedback activity cost at each time point is equal to the standard deviation of the parameter vector 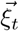 We computed the cost of feedback activity only at time points *t* when the optimal threshold values changed with respect to time point at *t* − 1. The resulting cost measure reflects the frequency of threshold switches and the range of parameter values which need to be transmitted from the observer to the sensory population via feedback connections after each switch.

### Inference tasks

#### Object detection

##### Environment dynamics and stimuli

At each trial, the environment switches randomly between two states corresponding to two values of the latent variable *θ*_*t*_: object present (*θ*_*t*_ = *P*) and object absent (*θ*_*t*_ = *A*), with the hazard rate *h* = 0.01. When the object was absent, stimuli *x*_*t*_ — samples from *p*(*x*_*t*_|*θ*_*t*_ = *A*) — were randomly drawn image patches with zero mean and unit variance. When the object was present, stimuli — samples from 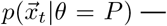 were a linear combination of a randomly selected image patch 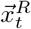, and pre-selected image of the object of interest 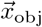 (a tree): 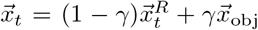, where the mixing coefficient *γ* = 0.2. Sparse coding neural activations *s*_*n,t*_ were determined using *λ* = 0.05 and 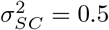

##### Observer model

At each time instant *t* the observer performed the following sequence of steps. First, the observer took the measurement *m*_*t*_ to be a projection of the image reconstructed from the sensory code 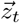 on the template image of the object of interest 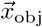, i.e., 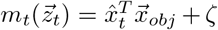 where *T* is vector transpose and *ζ* is a Gaussian noise with variance 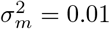.

Second, the observer updated the posterior distribution over the latent state *θ*:

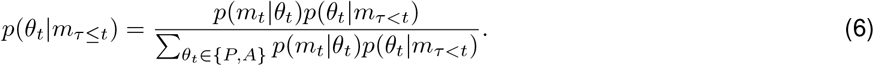

From the posterior, the observer computed the MAP estimate, 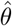. For simplicity, we assumed that 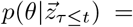
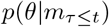 In the consecutive step, the observer computed the predictive distribution of the latent states 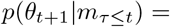 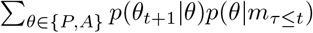. At low hazard rate, we could approximate that the predictive distribution is equal to the current posterior, *p*(*θ*_*t*+1_|*m*_*τ*≤*t*_) ≈ *p*(*θ*_*t*_|*m*_*τ*≤*t*_), from which we derived the predicted distribution of stimuli: 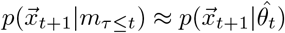

##### Nonlinearity optimization

To compute optimal nonlinearity thresholds for sensory encoding at different internal belief states of the observer, we first discretized the posterior distribution over the latent state into *k* = 32 bins, corresponding to linearly spaced values for *p*(*θ*_*t*_ = *P*|*m*_*τ*≤*t*_) over [0, 1]. Each of these states defined a distribution of expected image frames, 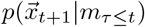. For each of these states, we generated a training dataset consisting of 10^4^ images with and without the object of interest mixed in proportion *p*(*θ*_*t*_ = *P*|*m*_*τ*≤*t*_)/(1 − *p*(*θ*_*t*_ = *P*|*m*_*τ*≤*t*_)). For each posterior state we then numerically optimized the Eq. (2) to derive optimal thresholds *ξ* at attentional resource constraint *ψ* = 4, using resilient-backpropagation gradient descent with numerically estimated gradient [62]. Each *ξ* was initialized with Gaussian noise. Since *ξ*_*n*_ ≥ 0, we performed the optimization with respect to real-valued auxiliary variables *a*_*n*_, where 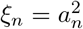. The resulting 32 vectors of optimal nonlinearity parameters 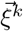 (where *k* ∈ {1, …, 32}) were used during subsequent simulations, where at each time step the observer selected the most appropriate set of nonlinearities *k*^*^:

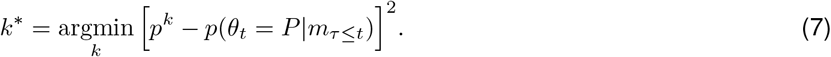

##### Simulation details

We generated a trajectory of the latent states of environment *θ*_*t*_ by concatenating 500 cycles of 50 samples of object present (*θ*_*t*_ = *P*) followed by 100 samples of object absent (*θ*_*t*_ = *A*) and again 50 samples of object present, resulting in the total length of 10^5^ time steps. Analyses in Fig. 4 B,C,E were performed by averaging over the 500 cycles. This artificial environment allowed us to compute averages over multiple changes of the latent state *θ*_*t*_.

#### Target localization

##### Environment dynamics and stimuli

The latent environmental state was defined by the 2D position of the center of the visual target (the white cross 7 × 7 pixels in size) 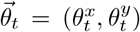, where *θ*_*x*_, *θ*_*y*_ ∈ {1, …, 32}. This position evolved as a random walk, 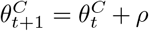, where 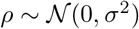 and *C* ∈ {*x*, *y*}; coordinates were rounded to nearest integer and bounded to image dimensions. We chose *σ* = 1.2 for the low-uncertainty scenario and *σ* = 2.4 for the high-uncertainty scenario to analyze the impact of uncertainty on the sensory code. The target was superposed on consecutive frames of a natural movie, 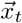. The presence of the artificially-designed visual target significantly changed statistics of images. Sparse coding neural activations *s*_*n,t*_ were determined using *λ* = 0.1 and 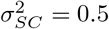.

##### Observer model

The observer computed the measurement 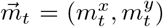 as the position of the peak of the two-dimensional cross-correlation function between the target template image (the cross) and the stimulus decoded from the neural code 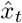. We assumed independent measurement noise in spatial coordinates for the measurement 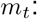 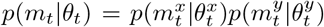, where marginal conditional distributions of coordinates are Gaussian: 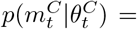 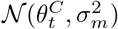(with *C* ∈ {*x, y*} is the index over spatial coordinates). To simplify optimization, we assumed vanishing measurement noise in this task, *σ*_*m*_ = 10^−5^.

The posterior distribution 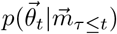 can be then computed separately for each spatial coordinate *C*:

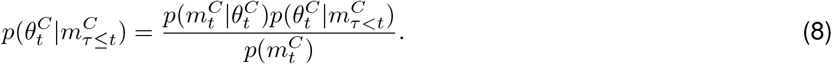

The prior distribution 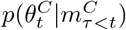 and the likelihood 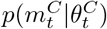 are Gaussian and conjugate to each other, therefore the posterior is also Gaussian, 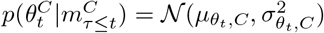 the point estimate for position is 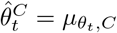

We further assume that the observer relies on trivial dynamics, where 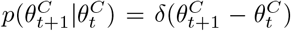 In this scenario, Eq (8) becomes a standard case of Bayesian online estimation of the mean with well-known closed form solutions [63].

Given these simpifying assumptions, the predicted distribution of measurements along each spatial coordinate is 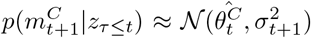 where the variance is the sum of the variance of the posterior and variance of the random walk i.e. 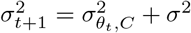

##### Nonlinearity optimization

We discretized the posterior belief about the position of the target into 25 values corresponding to a grid of 5 horizontal positions 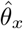 and 5 vertical positions 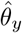 linearly spaced between 1 and 32 pixels. Nonlinearity optimization was performed analogously to the object detection task. At each time step the observer selected the optimal nonlinearity vector 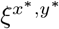 corresponding to the discretized position closest to the current position estimate 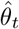:

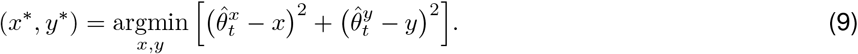

##### Simulation details

The simulation was ran for 10^4^ steps during which the target trajectory was evolving according to the dynamics described above.

#### Orientation estimation

##### Environment dynamics and stimuli

The environment state *θ*_*t*_ was switching randomly between two states with hazard rate *h* = 0.01. One of the states was generating images dominated by the vertical orientation *θ*_*t*_ = *V* and the other images with predominantly horizontal orientation *θ*_*t*_ = *H*. We identified these two states of the environment via unsupervised learning. First, we used the sparse coding model (without nonlinearities) to encode a large corpus of natural image patches 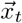. We then transformed activations of each model neuron *n* in response to each patch *t* by taking the log-ratio of its absolute value and the average magnitude of the activation of that neuron: 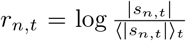 We then clustered such transformed vectors of the population response *r*_*t*_ into 9 clusters using the standard K-means algorithm. Out of these 9 clusters we visually selected two. One of these clusters included encodings of image patches where neurons with horizontally oriented basis functions were active stronger than their average. The other cluster included encodings of image patches where the vertically oriented basis functions were activated more strongly than the baseline. We selected these two sets of image patches to be generated by distributions 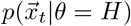 and 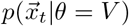 respectively. In this task, we used the following parameters of the sparse coding algorithm to encode the images: *λ* = 0.05 and 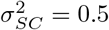.

##### Observer model

In this task the observer did not explicitly decode the image. Instead, it transformed neural activations *z*_*n,t*_ by taking their absolute value: *r*_*n,t*_ = |*z*_*n,t*_|. This vector of activity magnitude 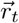 was then projected on the discriminative vector 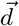 to obtain the measurement 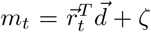, where *T* denotes vector transpose, and *ζ* is a Gaussian measurement noise with variance 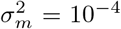. The discriminative vector 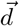 was a linear discriminant optimized to maximize discrimination accuracy between the two clusters of rescaled activity 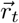 corresponding to the horizontal and vertical states respectively. We fitted distributions of noisy measurements *p*(*m*_*t*_ | *θ*_*t*_) with a Gaussian distribution for each state of the environment separately i.e. 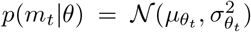, where *θ*_*t*_ ∈ {*V, H*}. The remaining computations were analogous to the object-detection task.

##### Nonlinearity optimization

Nonlinearity optimization was performed analogously to the object-detection task.

##### Simulation details

We generated a trajectory of the latent states of environment *θ*_*t*_ by concatenating 500 cycles of 50 samples of horizontal state (*θ*_*t*_ = *H*) followed by 100 samples of vertical state (*θ*_*t*_ = *V*) and again 50 samples of the horizontal state. Analyses in Fig. 4 B,C,E were performed by averaging over these 500 cycles.

### Computation of code statistics

#### Selection of task-modulated neurons

We sorted neurons according to how strongly they were modulated by the task. As a measure of the task modulation we took the ratio of the average activity of that neuron in the full sparse code and in the task-specific, adaptive code.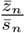 To compute activity correlation matrices in Fig. 5C, we selected 10 neurons with high modulation values computed in that way.

#### Response variability

To simulate response variability due to feedback modulation of the sensory code (Fig. 5D), we encoded the same, randomly selected image patch 1000 times while the belief of the observer was changing and adapting neural nonlinearities accordingly.

For the object detection and orientation estimation tasks we took the trajectory of the changing belief (*p*(*θ* = *P*) and *p*(*θ* = *H*) respectively) to be a sine function rescaled to fit in the interval [0.1, 0.9]. Over the 1000 stimulus presentations this sinusoid completed five cycles. For the target localization task we generated an instance of Gaussian walk, which determined the belief of the observer about the location of the target in the scene.

#### Noise correlations

For each task, we estimated noise correlations by computing correlation matrices of neural responses to 1000 presentations of the same stimulus (see above). To avoid numerical errors we added a Gaussian noise with variance *σ*^2^ = 0.01 to neural responses *z*_*n,t*_, after the stimulus has been encoded at each presentation. Correlations of the full code were all approximately equal to 0, since responses to each stimulus presentation were the same.

#### Code dimensionality, population activity and response fidelity as a function of perceptual uncertainty

To characterize the dimensionality of the code we computed PCA of the neural activity matrix *S*, where individual entries *s*_*n,t*_ are responses of the n-th neuron at t-th time point. We plotted the cumulative variance explained as a function of the number of principal components. For object detection and orientation estimation tasks we performed the dimensionality analysis by dividing the neural responses according to the level of uncertainty of the observer, and computing PCA on these responses separately. We quantified the uncertainty as the binary entropy of the prior over the latent state (*H*(*p*) = − *p* log_2_(*p*) − (1 − *p*) log_2_ (1 − *p*), where *p* is the probability of the object being present *p*(*θ* = *P*) in the object detection task, and the image orientation being horizontal *p*(*θ* = *H*) in the orientation estimation task. We defined three such intervals of uncertainty: [0, 0.33), [0.33, 0.66), and [0.66, 1] bits. For the target localization task, we run the simulation for two different levels of spatial uncertainty, determined by the variance of the target movements *σ*^2^.

To characterize the amount of population activity we computed the average absolute value of neural activations|*z*_*n,t*_|. The accuracy (or fidelity) of representation was computed as the average SNR dB of the image decoding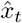 i.e 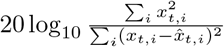, where *i* indexes the image pixels. For the object detection and orientation estimation tasks, we computed these average quantities for 10 levels of uncertainty spanned by the deciles of the uncertainty distribution. For the target localization task, we computed them for two different levels of spatial uncertainty, determined by the variance of the target movements *σ*^2^.

#### Determination of the number of active neurons

We declared *n*-th neuron to be active at time *t* if the magnitude of its activity exceeded the 1% of its maximal activity i.e. |*z*_*n,t*_| > 0.01 max_*t*_(|*z*_*n,t*_|). For each time point we computed the number of active neurons 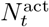, and averaged this number for different levels of uncertainty.

### Comparisons to data

#### Attentional modulation of population tuning curves

To estimate the population tuning curve, we first estimated orientation tuning curves of individual neurons. To do so, we fitted the basis function of each neuron with a parametric Gabor filter. In the next step, we parametrically varied the orientation of the Gabor filter, while keeping other parameters identical. We then discretized the interval of orientations 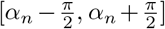 into 16 linearly spaced values, where *α*_*n*_ was the preferred orientation of that neuron. For each orientation value, we encoded the normalized image of the filter using the entire population, and took the neurons response *z*_*n*_(*ξ*_*n*_) to be a tuning at that orientation. We describe how we chose nonlinarity thresholds *ξ*_*n*_ below. We repeated this procedure for all neurons in the population. The population tuning curve was taken to be the average of tuning curves of individual neurons.

We ran a simulation of the target localization task for 10^4^ steps. The two population tuning curves in Fig. 6A were computed using different values of nonlinearity thresholds. To compute the population tuning curve in the absence of attention, for each neuron we took the nonlinearity threshold value averaged across the entire duration of simulation. To compute the population tuning curve in presence of attention, we took a single nonlinearity threshold value *ξ*_*n*_ corresponding to the belief that the target is closest to the spatial position of the Gabor filter encoded by that neuron.

#### Temporal statistics of gain dynamics

To compute temporal statistics of nonlinearity parameters we ran a simulation of the target localization task for 10^4^ steps. We note that while we computed temporal correlations of nonlinarity threshold parameters *ξ*_*n,t*_, the results do not qualitatively change if we take an inverse of the threshold 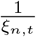 a parameter more directly related to the gain. As a measure of spatial tuning similiarity we took the correlation of the absolute values of neural basis functions |*ϕ*_*n*_|. We took the absolute value of neural nonlinearity outputs |*z*_*n,t*_| as a measure of neural activity level. Auto- and cross-correlation functions were computed using standard methods.

For the analysis displayed in Fig. 6 we selected only the neurons whose average activity magnitude ⟨|*z*_*n,t*_|⟩_*t*_ exceeded the 0.01 of the maximal activity average for all neurons in the population. The results do not qualitatively depend on this selection criterion.

## Acknowledgements

We thank Robbe Goris for generously providing figures from his work. W.M. was funded by the European Union’s Horizon 2020 research and innovation programme under the Marie Skłodowska-Curie Grant Agreement No. 754411.

## Supplemental Figures

**Figure S1:**
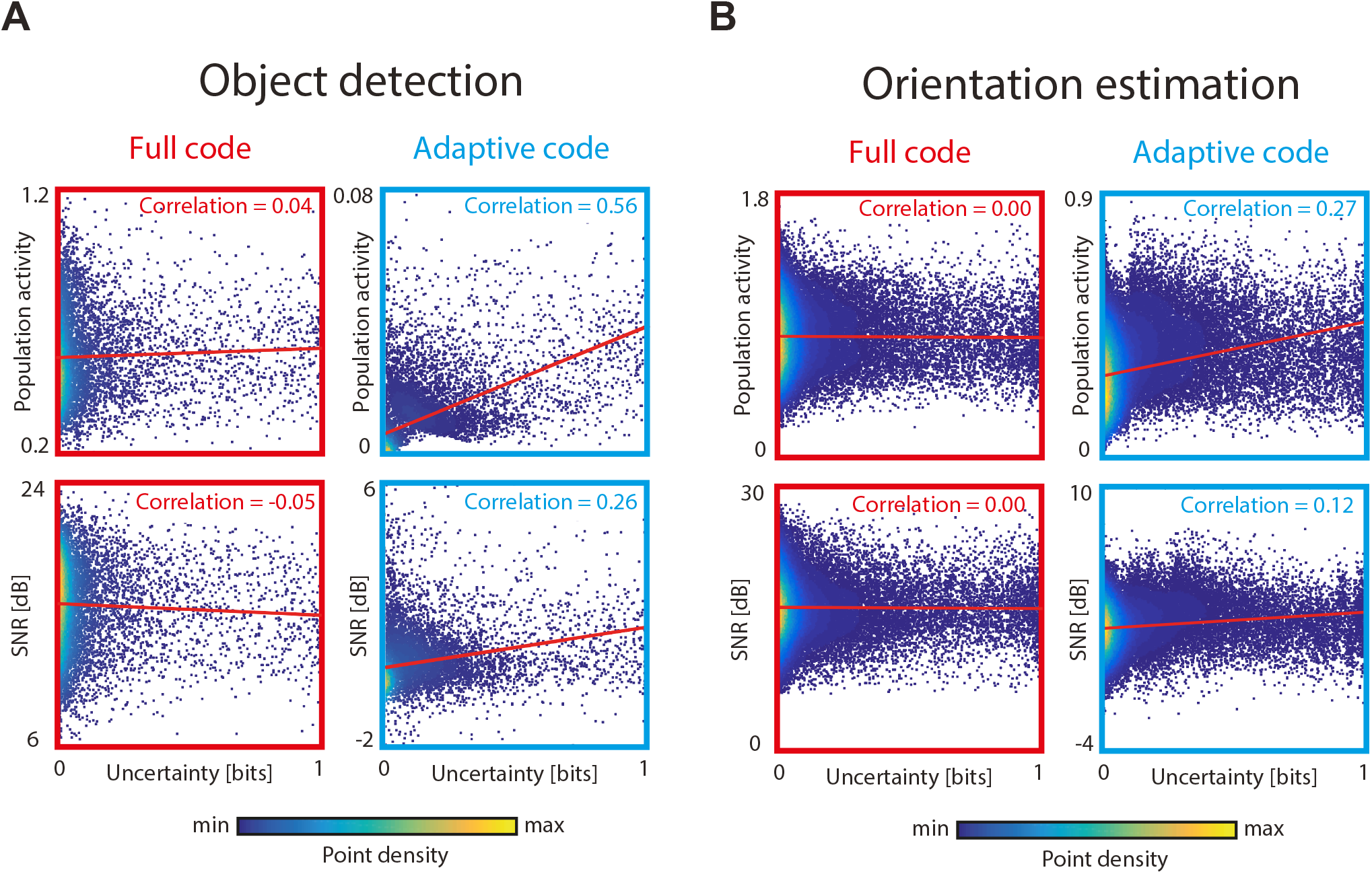
Statistics of uncertainty, population activity and representational fidelity. **A** Object detection task. Left column - full code (red) optimized for image reconstruction, right column (blue) adaptive code for inference. Top row - uncertainty vs population activity, bottom row - uncertainty vs representation fidelity. Each scatter density plot displays 10000 points. Red, dashed lines depict the linear fit. **B** Same as A but for the orientation estimation task.

**Figure S2:**
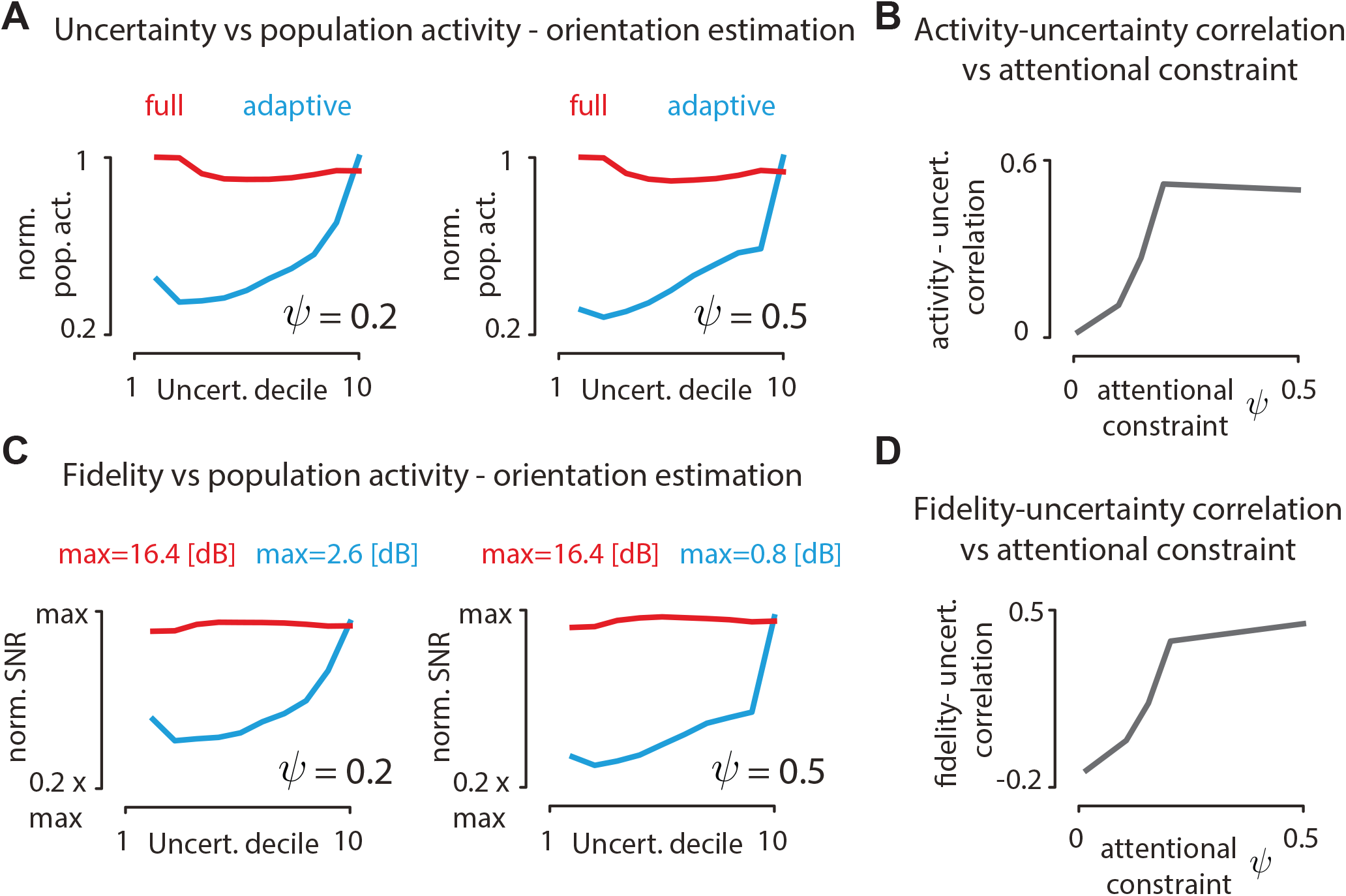
Impact of the attentional constraint *ψ* on uncertainty-activity and uncertainty-fidelity relations in the orientation-estimation task. **A** Uncertainty decile vs normalized population activity (analogous to Fig. 7B) for two values of the attentional constraint *ψ*. **B** Correlation between uncertainty and population activity as a function of the attentional constraint *ψ*. **C** Uncertainty decile vs encoding fidelity (analogous to Fig. 7D) for two values of the attentional constraint *ψ*. **D** Correlation between uncertainty and representation fidelity as a function of the attentional constraint *ψ*.

**Figure S3:**
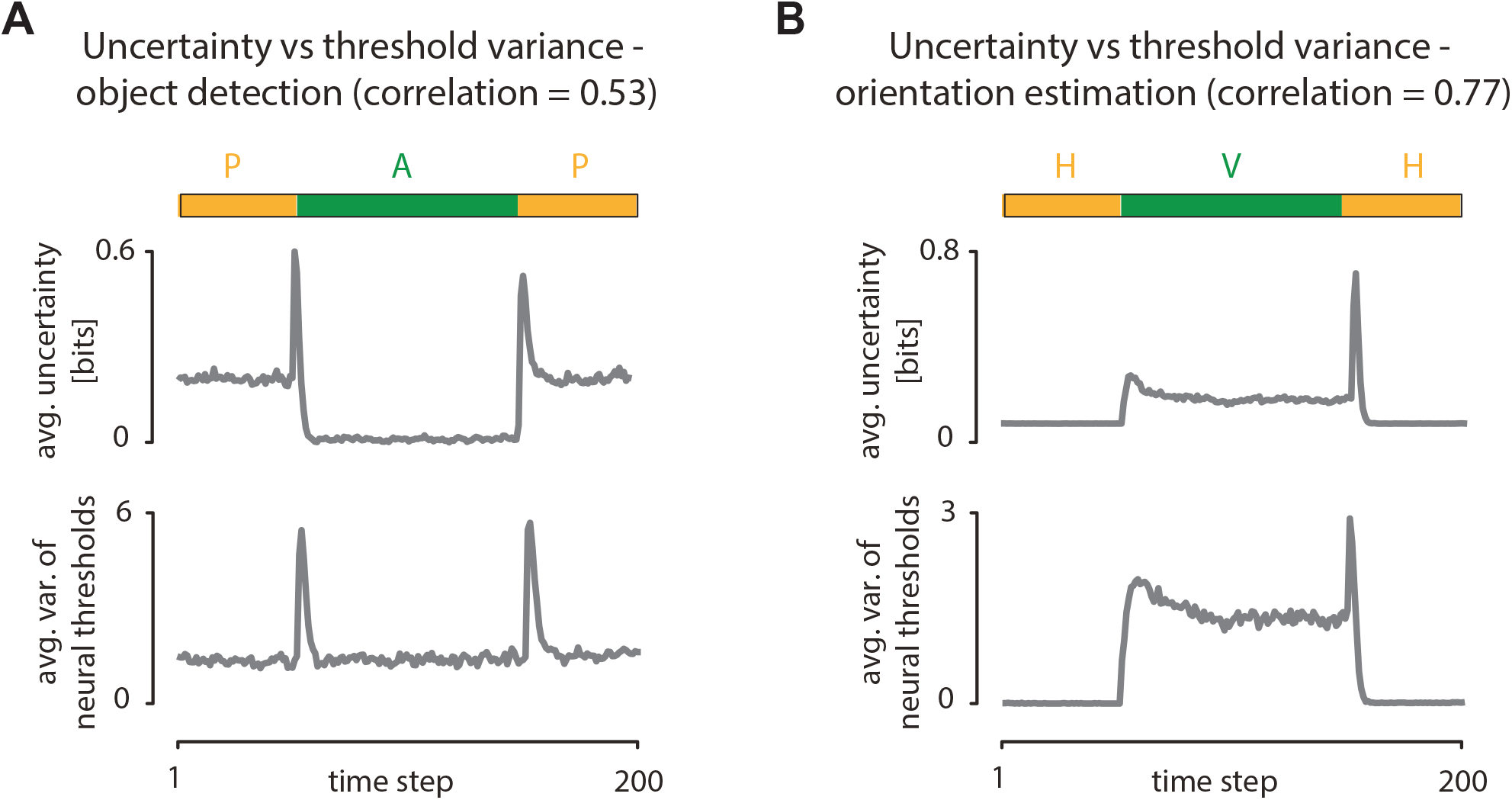
Average time courses of uncertainty and threshold (gain) variance. **A** Object detection task. Top - time course of posterior uncertainty (in bits) averaged over 500 switches between the environmental states (marked with a green-orange bar at the top). Bottom - time course of variances of neural thresholds *xi*_*n,t*_ averaged over 500 switches between the environmental states and neurons in the population. **B** Same as A but for the orientation estimation task.

